# LinearFold: Linear-Time Prediction of RNA Secondary Structures

**DOI:** 10.1101/263509

**Authors:** Dezhong Deng, Kai Zhao, David Hendrix, David H. Mathews, Liang Huang

**Author notes:** K.Z.’s contribution was done at School of EECS, Oregon State University. The authors declare no conflict of interest.

## Abstract

Predicting the secondary structure of an RNA sequence with speed and accuracy is useful in many applications such as drug design. The state-of-the-art predictors have a fundamental limitation: they have a run time that scales cubically with the length of the input sequence, which is slow for longer RNAs and limits the use of secondary structure prediction in genome-wide applications. To address this bottleneck, we designed the first linear-time algorithm for this problem. which can be used with both thermodynamic and machine-learned scoring functions. Our algorithm, like previous work, is based on dynamic programming (DP), but with two crucial differences: (a) we incrementally process the sequence in a left-to-right rather than in a bottom-up fashion, and (b) because of this incremental processing, we can further employ beam search pruning to ensure linear run time in practice (with the cost of exact search). Even though our search is approximate, surprisingly, it results in even higher overall accuracy on a diverse database of sequences with known structures. More interestingly, it leads to significantly more accurate predictions on the longest sequence families in that database (16S and 23S Ribosomal RNAs), as well as improved accuracies for long-range base pairs (500+ nucleotides apart).

---

Ribonucleic acid (RNA) is involved in numerous cellular processes. While many RNAs encode proteins (messenger RNAs, mRNAs), noncoding RNAs (ncRNAs) have intrinsic functions without being translated to proteins (1). ncRNA sequences catalyze reactions (2, 3), regulate gene expression (4–6), provide site recognition for proteins (7, 8) and serve in trafficking of proteins (9). The recent discovery and characterization of diverse classes of long noncoding RNAs, i.e. ncRNAs longer than 200*nt* (10), present new opportunities and challenges in determining their functions and mechanisms of action. Furthermore, the dual nature of RNA as both a genetic material and functional molecule led to the RNA World hypothesis, that RNA was the first molecule of life (11), and this dual nature has also been utilized to develop *in vitro* methods to evolve functional sequences (12). Finally, RNA is an important drug target and agent (13–18).

Predicting the secondary structure of an RNA sequence, defined as the set of all canonical base pairs (A–U. G–C, G–U). is an important and challenging problem (19, 20). Knowing structures reveals crucial information about the RNA’s function, which is useful in many applications ranging from ncRNA detection (21–23) to the design of oligonucleotides for knockdown of message (24, 25). Being able to rapidly determine the structure is useful given the overwhelming increase in genomic data (about 10^21^ base-pairs per year) (26) and given the small percentage of sequences that have experimentally determined structure. Experimental assays can provide information that can improve the accuracy of RNA secondary structure prediction (27), and these assays can now be used transcriptome-wide and *in vivo* (28–30). Recent studies focused on improved accuracy of prediction (31–35), but there is not enough attention on the speed of prediction.

**Fig. 1.**
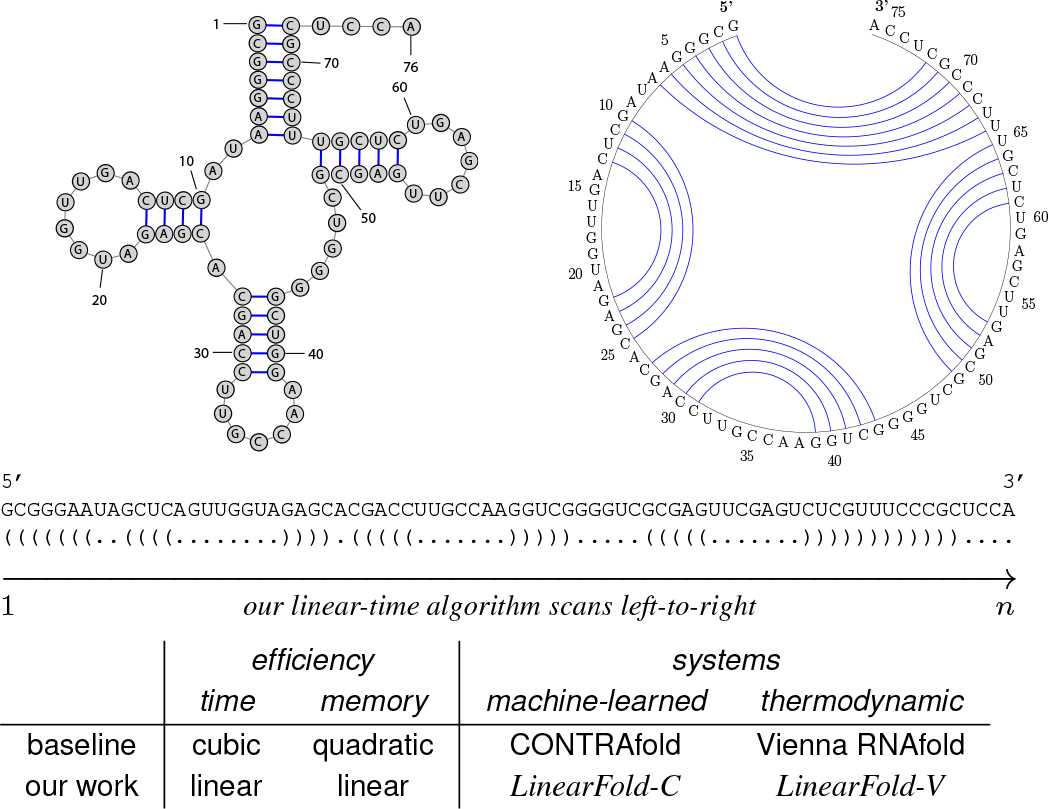
RNA secondary structure and high-level idea of our work. Top left: secondary structure of *E. coli* tRNA^Gly^; Top right: the corresponding circle plot; Central: the corresponding dot-bracket format. Bottom: schematic view of our work. In a nutshell, our algorithm scans the sequence left-to-right, and tags each nucleotide as “.” (unpaired), “(” (to be paired with a future nucleotide) or“)” (paired with a previous nucleotide).

### Significance Statement

Fast and accurate prediction of RNA secondary structures (the set of canonical base pairs) is an important problem, because RNA structures reveal crucial information about their functions. Existing approaches can reach a reasonable accuracy for relatively short RNAs but their running time scales almost cubically with sequence length, which is too slow for longer RNAs. We develop the first linear-time algorithm for RNA secondary structure prediction. Surprisingly, our algorithm not only runs much faster, but also leads to higher overall accuracy on a diverse set of RNA sequences with known structures, where the improvement is significant for long RNA families such as 16S and 23S Ribosomal RNAs. More interestingly, it also more accurate for long-range base pairs.

While there are two major approaches to *modeling* RNA secondary structures, namely the classical thermodynamic methods (36, 37) and the more recent machine learning-based methods (38, 39), all these efforts use virtually the same dynamic programming (DP) algorithm (40, 41) to find the best-scoring structure. However, this algorithm, borrowed from computational linguistics (42, 43), has a running time of *O*(*n*^3^) that scales *cubically* with the sequence length *n.* This is slow for long RNAs (*n* > 1,000), and in practice, many researchers resort to running this algorithm on short regions within the whole sequence, which inevitably ignores base pairs across segments (44). Computational and experimental studies demonstrate that base pairing between the ends of natural RNA sequences is expected.

In this paper, we design the first linear-time RNA secondary structure prediction algorithm, **LinearFold**, inspired by our previous work on linear-time natural language parsing (45). While the classical *O*(*n*^3^) dynamic programming is bottom-up, solving the best substructure for each span, our algorithm is left-to-right, incrementally tagging each nucleotide in the dot-bracket format (unpaired “.”, opening “(”, or closing “)”). While this naive version runs in exponential time *O*(3^*n*^), we use an efficient merging approach borrowed from computational linguistics (46) that reduces the running time back to *O*(*n*^3^). On top of this left-to-right algorithm, we further apply beam search, a popular heuristic to prune the search space (45), which keeps only the top *b* highest-scoring states at each nucleotide, resulting in an *O(n)* time approximate search algorithm. Even though our search is not exact, empirically, with a reasonable beam size (such as *b* = 100) it is close to exact search, and actually leads to better prediction accuracies than exact search.

Our algorithm can be used with both thermodynamic and machine learned models. In particular, we implemented two versions of LinearFold, *LinearFold-V* using the thermodynamic free energy model from Vienna RNAfold (37), and *LinearFold-C* using the machine learned model from CONTRAfold (38) (see Fig. 1 (bottom)). We evaluate our systems on a diverse dataset of RNA sequences with well-established structures, and show that while being substantially more efficient, *LinearFold* leads to higher average accuracies over all families, and somewhat surprisingly, *LinearFold* is significantly more accurate than the exact search methods on the longest families 16S and 23S Ribosomal RNAs. More interestingly, *LinearFold* is also more accurate on long-range base pairs that are more than 500 nucleotides apart, which is well known to be a challenging problem for the current models (47).

## Results

### Efficiency and Scalability of LinearFold

To demonstrate the efficiency and scalability of *LinearFold*, we compare its running time with the conventional cubic-time prediction algorithms used in the baseline systems, CONTRAfold and Vienna RNAfold. Figure 2 shows the results on two datasets: (a) the ArchiveII dataset (48), a diverse set of RNA sequences with known structures (see details in the Methods section and Table SI 1), and (b) a (sampled) subset of RNAcentral (49), a comprehensive (meta-)set of ncRNA sequences from many databases. While the ArchiveII set contains sequences of length 3,000 or less, the RNAcentral set has many much longer sequences, with the longest being 244,296 *nt* (Homo Sapiens Transcript NONHSAT168677.1, from the NONCODE database (50)). We use a machine with 3.40GHz Intel Xeon CPUs and 32G memory, running Linux; all programs are written in C/C++ compiled by GCC 4.9.0.

**Fig. 2.**
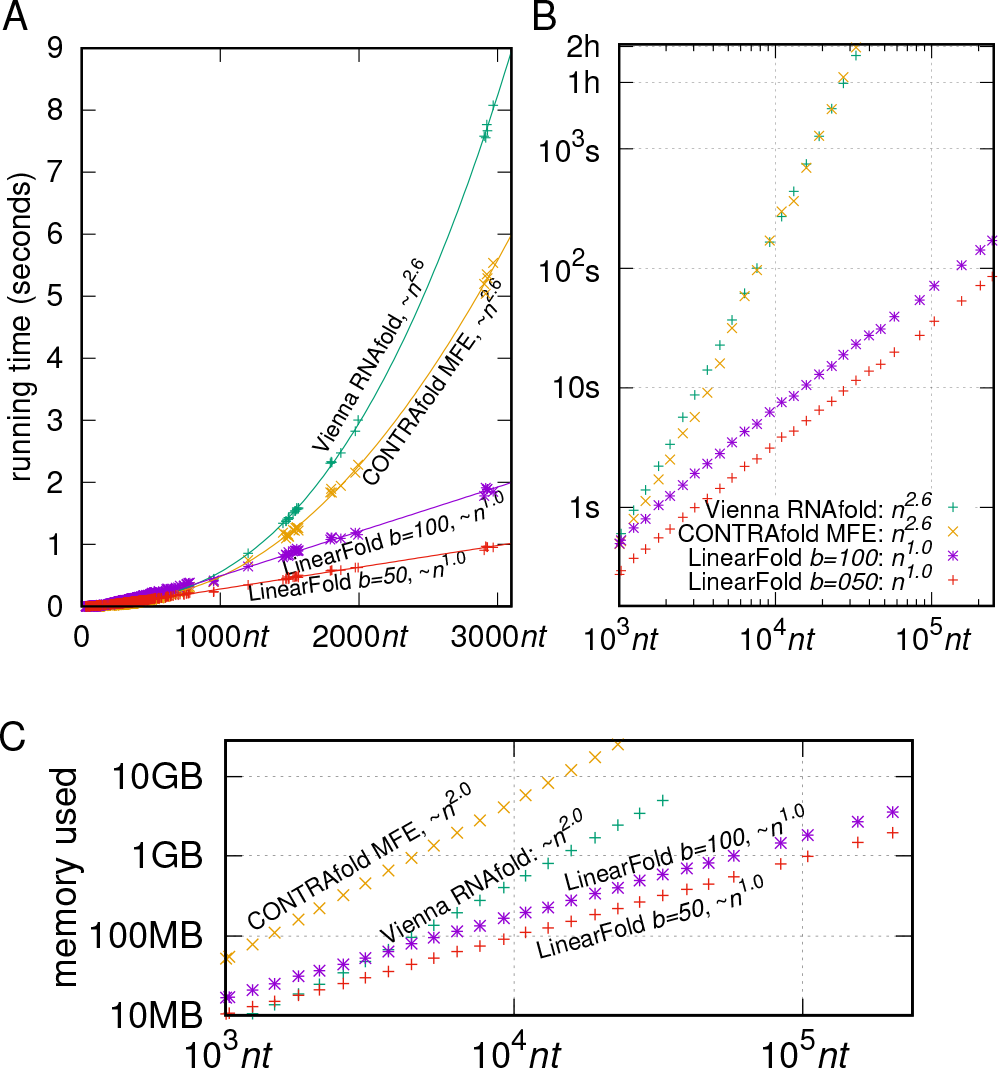
A: runtime comparisons on ArchiveII dataset: *LinearFold-C* (with beam sizes 100 and 50) vs. two baselines, CONTRAfold MFE & Vienna RNAfold (*LinearFold-V* have identical running time with *LinearFold-C).* B: runtime comparisons on RNAcentral dataset (log-log). C: memory usage comparisons (RNAcentral set, log-log).

Figure 2 A confirms that *LinearFold*’s running time scales linearly with the sequence length, while the two baseline systems scale super-linearly, with an empirical runtime of *O*(*n*^2.6^) determined by curve fitting. Figure 2 B reconfirms this fact on much longer sequences (in log-scale), and for a sequence of ~10,000 *nt* (e.g., the HIV genome), *LinearFold* (with the default beam size of *b*=100) takes only 7 seconds while the baselines take 4 minutes. For sequences of length 32,753, our *LinearFold* takes only 23 seconds while CONTRAfold takes 2 hours and RNAfold 1.7 hours. This clearly shows the advantage of our linear-time prediction algorithm on very long ncRNAs.

In addition, *LinearFold* also has an advantage on memory usage that leads to better scalability on extremely long sequences. The baseline cubic-time algorithms require memory space that scales *quadratically* with sequence length, because intuitively they need to figure out the best scoring or minimum free energy structure for every substring *[i,j]* of the entire sequence. This means you need 4x memory if your sequence length doubles. In addition, due to a design deficiency, neither CONTRAfold MFE or Vienna RNAfold runs on any sequence longer than 32,767 *nt*. On the other hand, *LinearFold* not only takes linear time, but also uses linear memory, without the need for the two-dimensional table of size *O*(*n*^2^). As a result, *LinearFold* is able to process the longest sequence in RNAcentral (244,296 *nt),* taking less than 3 minutes. In fact, *LinearFold* even scales to sequences of 10,000,000 *nt* on our 32GB-memory machine.

### Accuracy of LinearFold

We next compare the prediction accuracies of *LinearFold* and the two baseline systems, reporting both Positive Predictive Value (PPV; the fraction of predicted pairs in the known structure) and Sensitivity (the fraction of known pairs predicted) on each RNA family in ArchiveII dataset. We also tested statistical significance using a paired, two-tailed *t*-test, following previous work (51). Figure 3 shows that *LinearFold-C* improves PPV and Sensitivity over CONTRAfold by +2.1%/+1.4% (absolute)when averaged across all families. This is surprising because *LinearFold* produces more accurate structures using a fraction of runtime. Individually, *LinearFold-C* is significantly more accurate in both PPV/Sensitivity on three families: Group I Intron, 16S and 23S ribosomal RNAs, with the last two being the longest families in this dataset. 16S rRNAs have an average length of 1548 *nt* and +3.89%/+3.08% absolute improvement in PPV/Sensitivity, and 23S rRNAs have an average length of 2927 *nt* and +14.00%/+9.98% absolute improvement. Even more surprising, *LinearFold*’s improvement in accuracy is more pronounced on longer sequences. Accuracies are also improved by *LinearFold-V* over Vienna RNAfold, but the difference is smaller (overall +0.2%/+0.2% absolute improvement in PPV/Sensitivity). Individually, the improvement is significant on both PPV/Sensitivity on two families, Group I Intron and 16S rRNA.

**Fig. 3.**
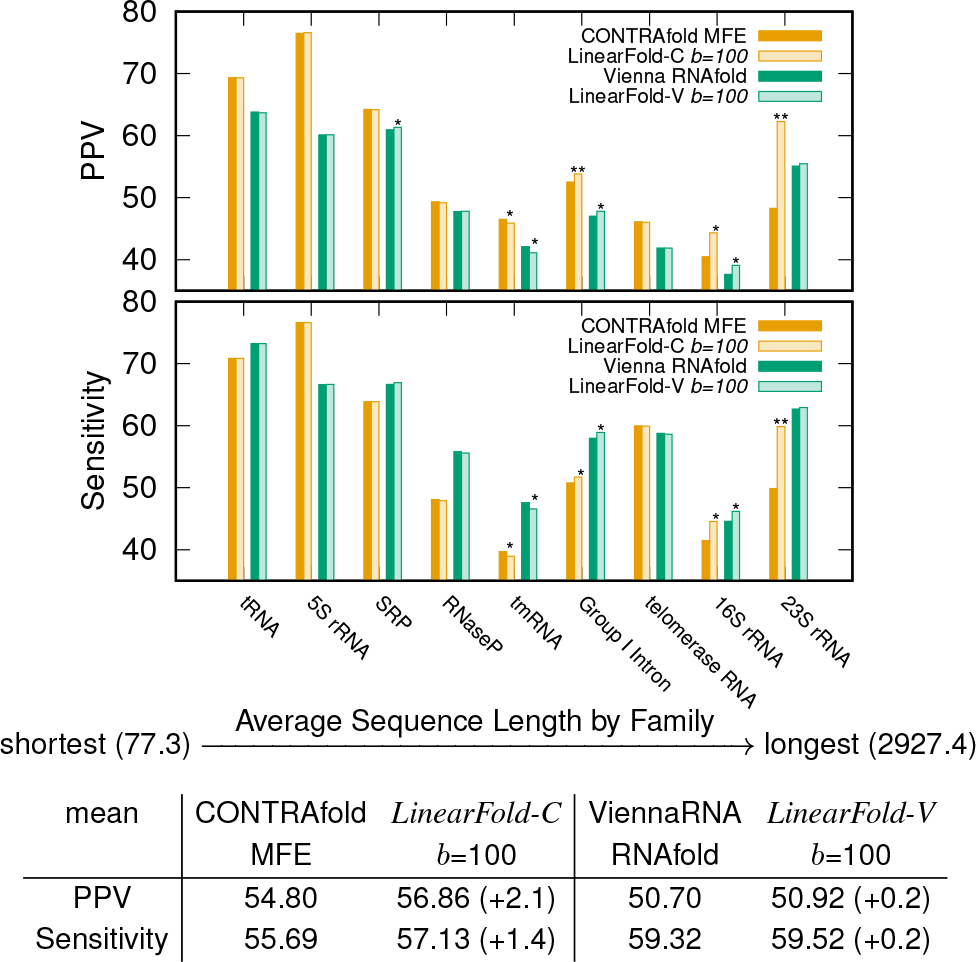
PPV and Sensitivity (by family) on the ArchiveII dataset, comparing *LinearFold* with the corresponding baselines, CONTRAfold MFE and Vienna RNAfold. Each column represents a family accuracy, averaged over all sequences in that family. The overall accuracies are reported in the table, averaging over all families. Statistical significance is marked as *(0.01 < *p* < 0.05), and *\*\*{p* < 0.01). See Table SI 1 for detailed accuracy numbers. See the Methods section for details of the PPV/Sensitivity metrics and the significance testing method.

### Impact of Beam Size

Above we used *b*=100 as the default beam size. Now we investigate the impact of different beam sizes. We first study the impact of search quality. Since our *LinearFold* algorithm uses approximate search instead of exact search, we use the difference between exact search free energy and our returned free energy as the measure of search quality – the closer they are, the better our search quality. We can see from Figure 4 A that the search quality is getting closer when the beam size increases, and *Linear-Fold* achieves similar model cost / free energy using the default beam size. Similarly, Figure 4 B plots the number of pairs predicted, at each beam size. It shows that ViennaRNA model tends to overpredict, while the CONTRAfold model underpredicts. Also, although our algorithm always underpredicts compared to exact predictions in either model, we predict almost the same number of base pairs as each model when using the default beam size. The mean difference is 0. 29(CONTRAfold)/0.01 (ViennaRNA) pairs when *b =* 100.

**Fig. 4.**
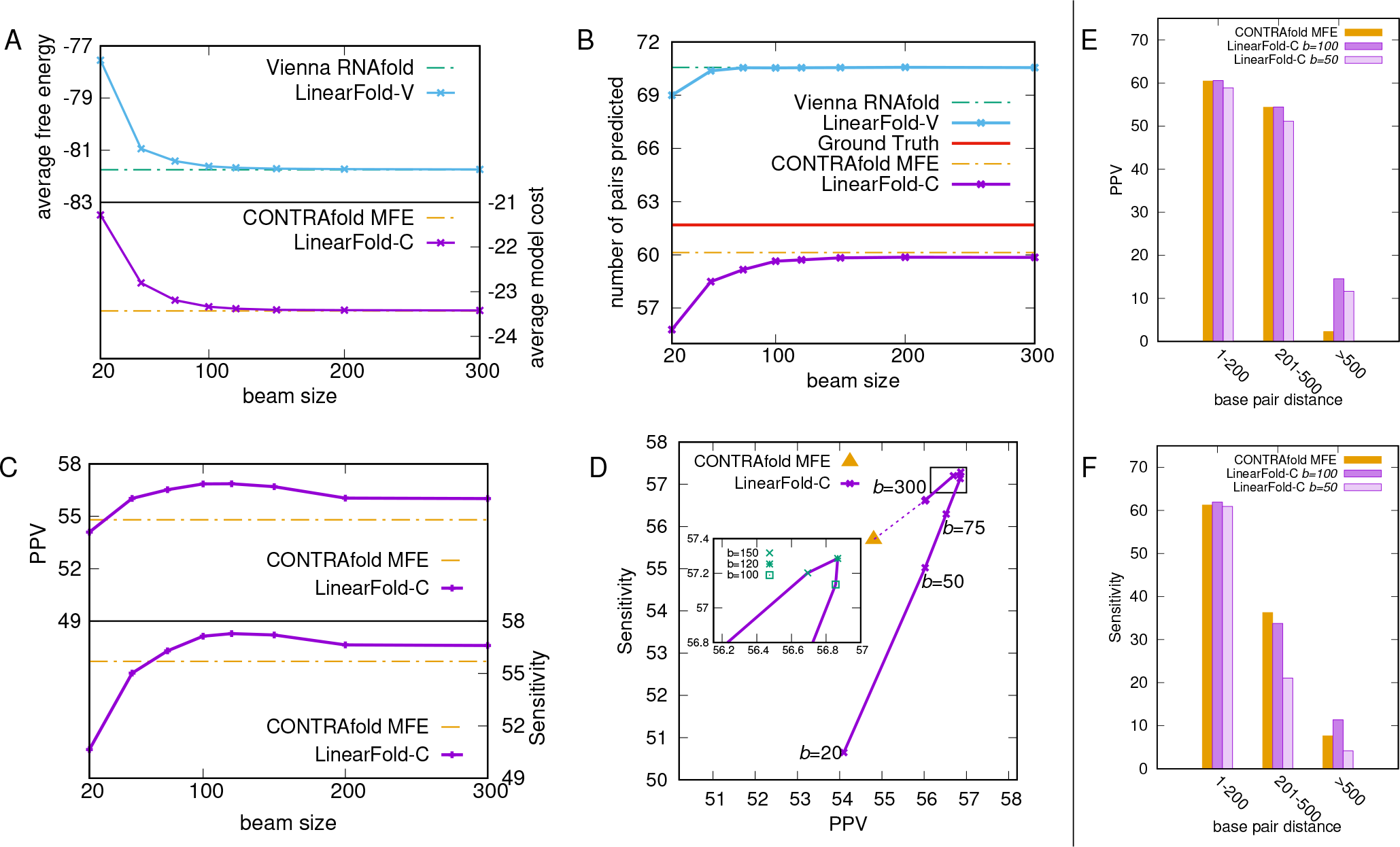
Impact of beam size. A-D illustrate the trends of different variables when beam size increases. A: the internal cost, namely averaged free energy / model cost of two versions of *LinearFold*; B: the number of pairs predicted (averaged by sequence) of these methods, comparing with Ground Truth; C: change of both PPV and Senstivity with the increasing of beam size; D: PPV-Sensitivity tradeoff when varying beam size; E-F: PPV and Sensitivity against pair distance in the ArchiveII dataset, comparing *LinearFold-C* with CONTRAfold MFE. Each point represents the overall PPV/Sensitivity of all base pairs in a certain length range.

Figure 4 C plots PPV and Sensitivity as a function of beam size *b. LinearFold-C* outperforms CONTRAfold MFE in both PPV and Sensitivity with *b* ≥75 (though it will converge to CONTRAfold MFE when *b* ⟶ +∞). and *LinearFold-C’s* PPV/Sensitivity are stable with *b* ∈ [100,150]. Figure 4 D shows the tradeoff of PPV and Sensitivity of *LinearFold*, with the change of the beam size. It starts with the increasing of both PPV and Sensitivity, reaches the peak at *b*=120, and falls until converging to exact search. Although the peak happens when beam size is 120, we can see that the performance *LinearFold* is consistent when its beam size is in [100, 150], as both PPV/Sensitivity stays almost the same.

### Accuracy Improvements on Long-Range Base Pairs

We further evaluated the effect of the distance between base paired-nucleotides on prediction performance. As shown in Figure 4 E and F, when predicting long-distance pairs, *LinearFold* can outperform previous approaches in both PPV and Sensitivity. Contrary to the concern that *LinearFold* would not predict as accurately for long-distance pairs, it continues to outperform previous methods even at pairing distances over 500 nt. Detailed comparisons between *LinearFold-V* and Vienna RNAfold in PPV, Sensitivity, and prediction quality, are in the Supporting Information.

### Example Predictions: Group I Intron, 16S and 23S rRNAs

We visualized the predicted secondary structure of 3 examples from different RNA families, Group I Intron C. *Mirabilis*, 16S rRNA *A. Pyrophilus,* 23S rRNA *E. coli,* comparing *LinearFold-C* with CONTRAfold MFE (Figure 5). The circular plots show our performance improvement, by both predicting more base pairs correctly, and reducing incorrect predictions. This visualization also demonstrates *LinearFold’s* improved prediction of long-distance base pairs than the baseline, as shown in Group I Mirabilis (bottom half, pair distance 250), 16S *A. Pyrophilus* (left part, 550), 23S *E. coli* (left part, 600). Fig. SI 3 shows the corresponding results comparing *LinearFold-V* with Vienna RNAfold. We also built a web demo visualizing results from all sequences in these three families.^*^

**Fig. 5.**
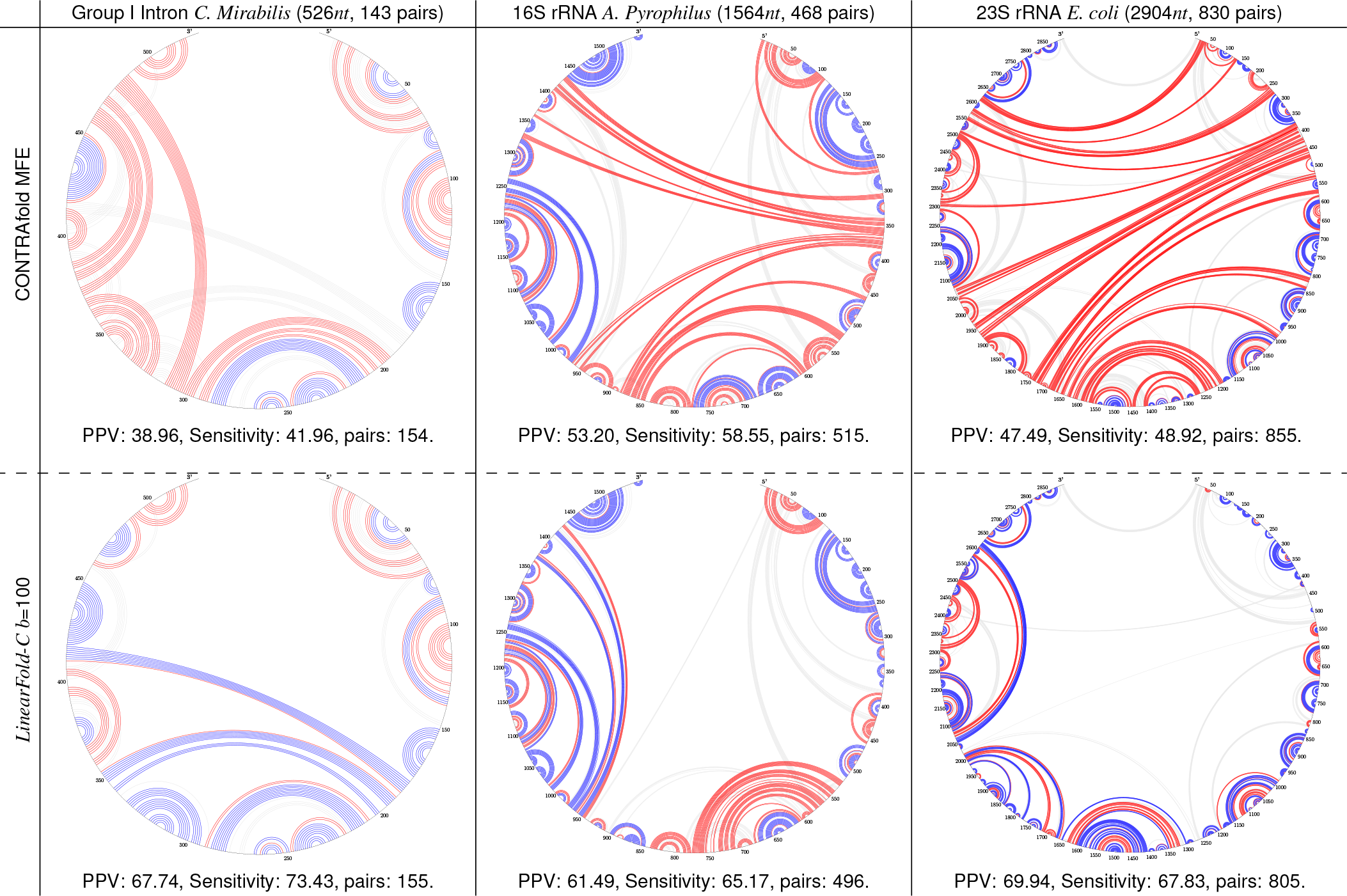
Circular plots of 3 RNA sequences (selected from 3 different RNA families) comparing CONTRAfold MFE with *LinearFold-C.* A blue arc represents a correctly predicted pair, a red arc represents an incorrectly predicted pair, while a light gray arc represents a based pair miss predicted. Each plot uses the clockwise order of the RNA sequence from the top. These 3 examples are picked from 3 different RNA families that the performance of *LinearFold-C* is improved significantly comparing to CONTRAfold MFE.

## Discussion

RNA structure prediction is important for inferring RNA function and has many applications including drug design. The existing algorithms for RNA secondary structure prediction run in time that scales cubically with the sequence length, which is too slow for long non-coding RNAs; e.g., the baseline systems in this work, CONTRAfold and Vienna RNAfold, which are two of the most popular prediction software, take 2 hours and 1.7 hours, respectively, for a sequence of 32,753 *nt*. Furthermore, the existing algorithms also need memory space that scales quadratically with the sequence length, and as a result, both baseline systems fail to run on sequences beyond 32,767 *nt*. In reality, the longest RNA sequence in the RNAcentral dataset is 244,296 *nt*, which is 7× of that limit.

We design the first linear-time, linear-memory prediction algorithm, *LinearFold*, using dynamic programming plus beam search, and apply this algorithm to both machine-learned and thermodynamic models. The linearity in both time and memory is confirmed in Fig. 2. What is more interesting is the following three surprising findings:

1. 1. Even though *LinearFold* uses only a fraction of runtime and memory compared to the existing algorithms in the baseline systems, our predicted structures are overall more accurate in both PPV and Sensitivity and on both machine-learned and thermodynamic models (see Fig. 3).
2. 2. The accuracy improvement of *LinearFold* is more pronunced on longer families such as 16S and 23S rRNAs (see Figs. 3 and 5).
3. 3. Even more surprisingly, *LinearFold* is also more accurate than the baselines on long-range base pairs that are more than 500 *nt* apart (Fig. 4 E–F), which is well known to be a challenging problem for the current models (47).
4. 4. Although the model depends on the beam size *b*, the number of base pairs and the accuracy of prediction are very robust to changes in beam size (when *b* is in the range of 100-200).

Why our beam search algorithm, even though being approximate, outperforms the exact search baselines in terms of accuracy (esp. in 16S and 23S rRNAs)? First, current thermodynamic and machine learned models are far from perfect, so it is totally possible that a suboptimal structure (in terms of free energy or model score) is more accurate (in terms of PPV/Sensitivity) than the optimal structure. For example, for sequences of about 400 nucleotides, a structure about 80% correct can be found with a free energy within 5% of the optimal structure (52). But how does our algorithm *systematically* pick a more accurate suboptimal structure without seeing the ground truth? We suspect that it is because beam search prunes lower-scoring (sub)structures at each step, requiring the surviving (sub)structures to be highly scored at each prefix. This extra constraint might compensate for the inaccuracy of the model.

Our algorithm has several potential extensions. First of all, it might be possible to extend *LinearFold* to calculate the partition function and base pair probabilities, since the classical method for that task, the McCaskill algorithm (53), is similar to the cubic-time structure prediction algorithms which are used as baselines in this paper. Secondly, this linear-time algorithm to calculate base pair probabilities should facilitate the linear-time identification of pseudoknots, by replacing the cubic-time McCaskill algorithm with a linear-time one in those pseudoknot-prediction programs (54, 55). Thirdly, being linear-time, *LinearFold* also facilitate easier and faster training of parameters than the cubic-time CONTRAfold using structured prediction methods (56), and we envision a retrained model tailored to linear-time prediction should be even more accurate.

## Methods

Our *LinearFold* approach is presented in four steps, starting from the most naive but easy-to-understand exhaustive search version (Fig. 6 A), and gradually build it up to the linear-time version (Fig. 6 D), using a graph-structured stack and beam search.

**Fig. 6.**
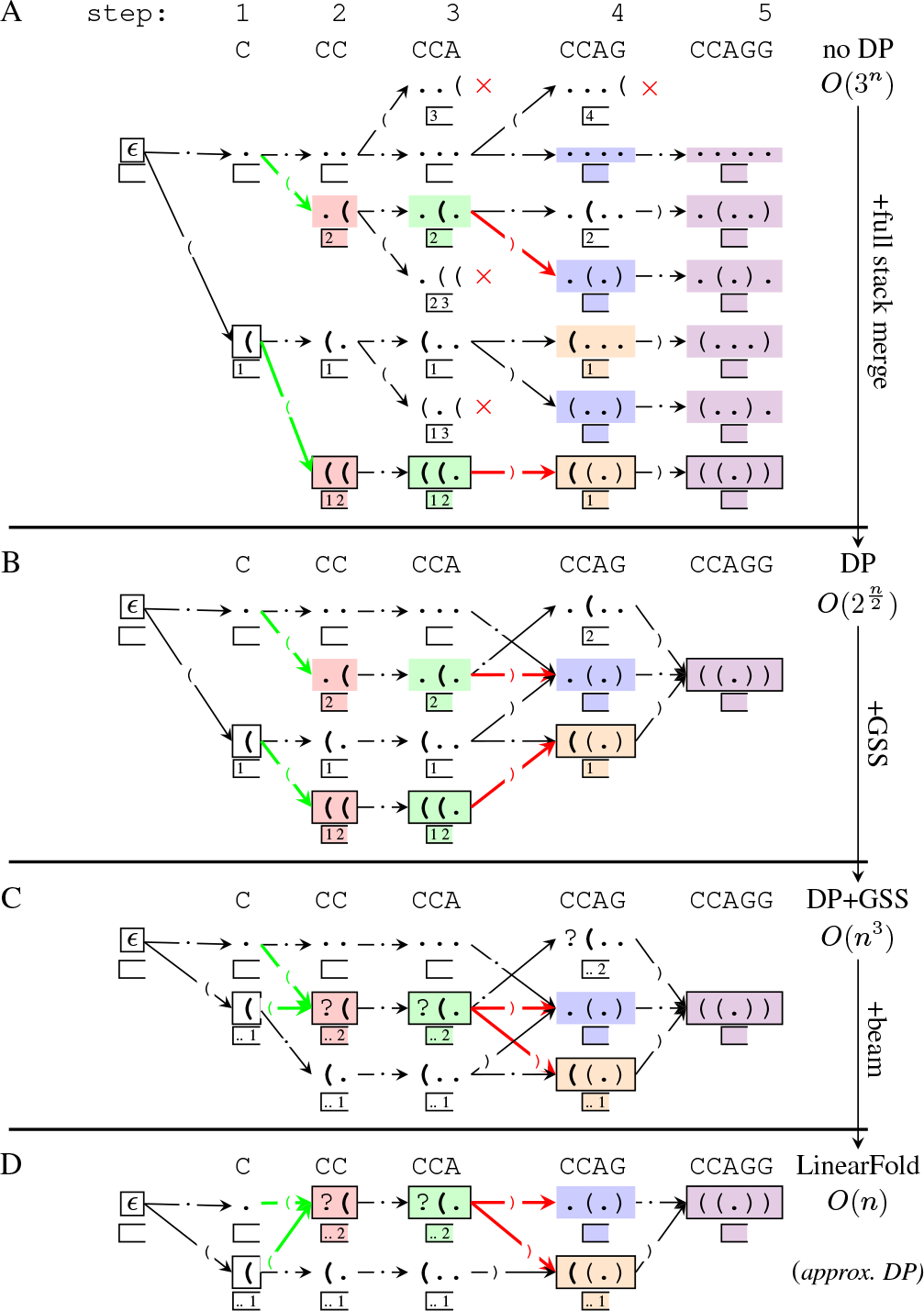
Four-step demonstration of LinearFold, simply finding the max number of pairs instead of the actual MFE model. Each node represents a predicted prefix of the structure, with a stack showing the unpaired openings; each arrow corresponds to an action (push, skip, and pop); dead-end states are with a red ×. In C, nodes with the same color are merged by the Graph-Structured Stack. Nodes with borders represent the ground truth path.

The basic idea of linear-time prediction is to predict incrementally from left to right, labeling each nucleotide as unpaired “•”, opening “(”, or closing “)” We require this dot-bracket string to be well-balanced as we only consider pseudoknot-free structures.

Given an input RNA sequence x = *x*_0_*x*_1_ … *x_n_-i* where *x_i_* ∈ {A, C, G, U}, our algorithm aims to find the best structure y = *y*_0_ *y*_1_ … *y*_*n*_–1 where *y_i_* {“.”, “(”, “)”} with minimum free energy (or minimum model cost):

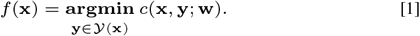

Here *Y*(x) is the set of all possible structures, i.e., {y | y has balanced parentheses}, *c* is the cost function (i.e., free energy function), and w is the model (and parameters).

**Naive exhaustive incremental prediction:** *O*(3^*n*^) **time.** By exhaustively predicting *y* from left-to-right, we traverse all the possible structures in *y*(x), and pick the one with the minimum free energy or model cost. We formalize each state at step *j* (*j* ∈ {0, …, *n*}) to be a triple, *s* = ⟨σ|*i, j*⟩: *y*, where σ|i is a stack consisting of unmatched openings so far where *i* is the top of the stack, meaning *x_i_* is the last unmatched opening nucleotide. *y* is the corresponding dot-bracket (sub)sequence up to *x_j_* – *i*. For each state, it can transition to a subsequent state, taking one of the three actions: push, which labels the current nucleotide *x_j_* as a left bracket “(”, putting it on top of the stack, skip, which labels *x_j_* as a dot “.” leaving the stack unchanged, and pop, which labels *x_j_* as a right bracket “)”, if it matches *x_i_* and popping *i* from the stack. See Fig. SI 4 (a) for the deductive system. This algorithm takes *O*(3^*n*^) time to exhaustively traverse all possible structures (see Figure 6 A).

**Dynamic Programming via Full Stack Merging:** 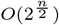 **time.** Now we apply dynamic programming on top of this exhaustive method to exploit shared computations. Consider a simple case that two states can be merged: if there are two states in the same step *j,* ⟨σ, *j*⟩: *y’* and ⟨σ, *j*⟩: *y’* sharing the exact same stack σ but with different dot-bracket strings *y* and *y’*, we say that these two states are “equivalent” and we can merge them (and only keep the better scoring between *y* and *y’*. Fig. 6 B illustrates this merging. Although we merge to reduce the number of states, it is still exponential time, since there could be exponentially many different stacks in each step. This algorithm takes 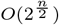 time.

**Dynamic Programming via Graph Structured Stacks:** *O*(*n*^3^) **time.** To avoid considering exponentially many states, we further merge states with different stacks. Consider two states in the same step *j*, ⟨σ_0_|*i*, *j*⟩ and ⟨σ_1_|*i*, *j*⟩ which share the last unpaired opening *i* (i.e., stack top). We call these states “temporarily equivalent”, since they can be treated as exactly the same until the unpaired opening *x_i_* is closed (and thus popped from the stack). In other words we can represent both stacks σ_0_|*i* and σ_1_|*i* as … *i* where … denotes part of the history that we do not care at this moment. This factorization of stacks is called “Graph-Structured Stacks” (GSS) by Tomita (46). After merging, we define the new state to be ⟨*i, j*⟩ and therefore we maintain *O*(*n*^2^) states. For each state ⟨*i, j*⟩, the pop action can take worst-case *O*(*n*) time because ⟨*i, j*⟩ can combine with every ⟨*k, i*⟩ from step *i*. Thus the overall time complexity is *O*(*n*^3^). See Fig. 6 C for an example of the merging process and Fig. SI 4 (b) for the deductive system.

**Dynamic Programming via Beam Search:** *O(n)* **time.** In practice, the exact search algorithm still runs in *O*(*n*^3^) time. But this left-to-right *O*(*n*^3^) search is easily “linearizable” unlike the traditional bottom-up *O*(*n*^3^) search used by all existing systems for RNA structure prediction. We further employ beam search pruning (56) to reduce the complexity to linear time. Generally, we only keep the *b* top-scoring states ⟨*i, j*⟩ for each step. This way all the lower-scoring states are pruned, and if a structure survives to the end, it must have been one of the top *b* states in every step. This pruning also means that in a pop action, a state (*i, j*) can combine with at most *b* states (*k, i*) from step *i*. Thus the overall time complexity is *O*(*nb*^2^). However, instead of generating *b*^2^ new states from a pop action, we use **cube pruning** (57) to generate the best *b* states, which would take *O*(*b* log *b*) time. Thus the overall running time over a length-*n* sequence is *O*(*nb* log *b*), see See Figure 6 D for beam search.

**Dataset, Evaluation Metrics and Significance Testing.** We choose the ArchiveII dataset (48), a diverse set of over 3,000 RNA sequences with known secondary structures. But since the current CONTRAfold machine-learned model (v2.02) is trained on the S-Processed dataset (58) we removed those sequences appeared in the S-Processed dataset. The resulting dataset we used contains 2,889 sequences over 9 families, with an average length of 222.2 *nt*. Due to the uncertainty of base-pair matches existing in comparative analysis, we consider a base pair to be correctly predicted if it is also slipped by one nucleotide on a strand, accordingly((48)). Generally, if a pair (*i, j*) is in the predicted structure, we claim it’s correct if one of (*i, j*), (*i* – 1,*j*), (*i* + 1,*j*), (*i,j* – 1), (*i,j* + 1) is in the ground truth structure. We report both Sensitivity and PPV where

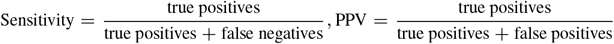

We use the paired two-tailed *t*-test to calculate the statistical significance, with the type I error rate, consistent with the previous methods (51).

## ACKNOWLEDGMENTS

This work was partially supported by NSF grant 1656051 (L.H.), NIH grants R56 AG053460 and R21 AG052950 (D.H.), and NIH grant R01GM076485 (D.H.M.). We thank James Cross for algorithm design, Kaibo Liu for the web demo, and Juneki Hong for proofreading.

## Supporting Information

**Table SI 1.**
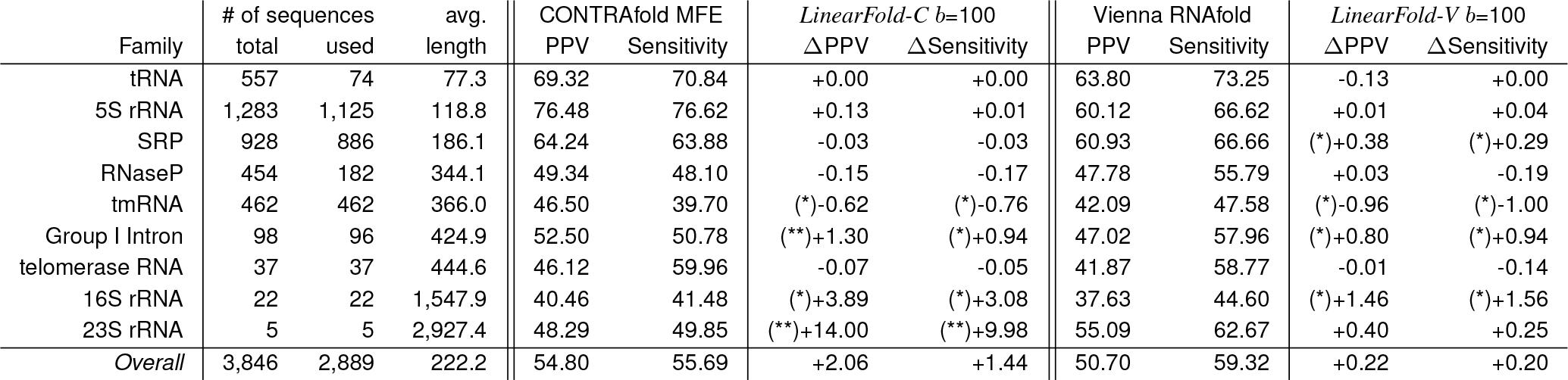
Detailed information of the ArchiveII dataset and the prediction accuracies of CONTRAfold MFE, *LinearFold-C*, Vienna RNAfold and *LinearFold-V*. Statistical significance are marked by *(0.01 ≤ *p* < 0.05) and **(*p* < 0.01).

**Fig. SI 1.**
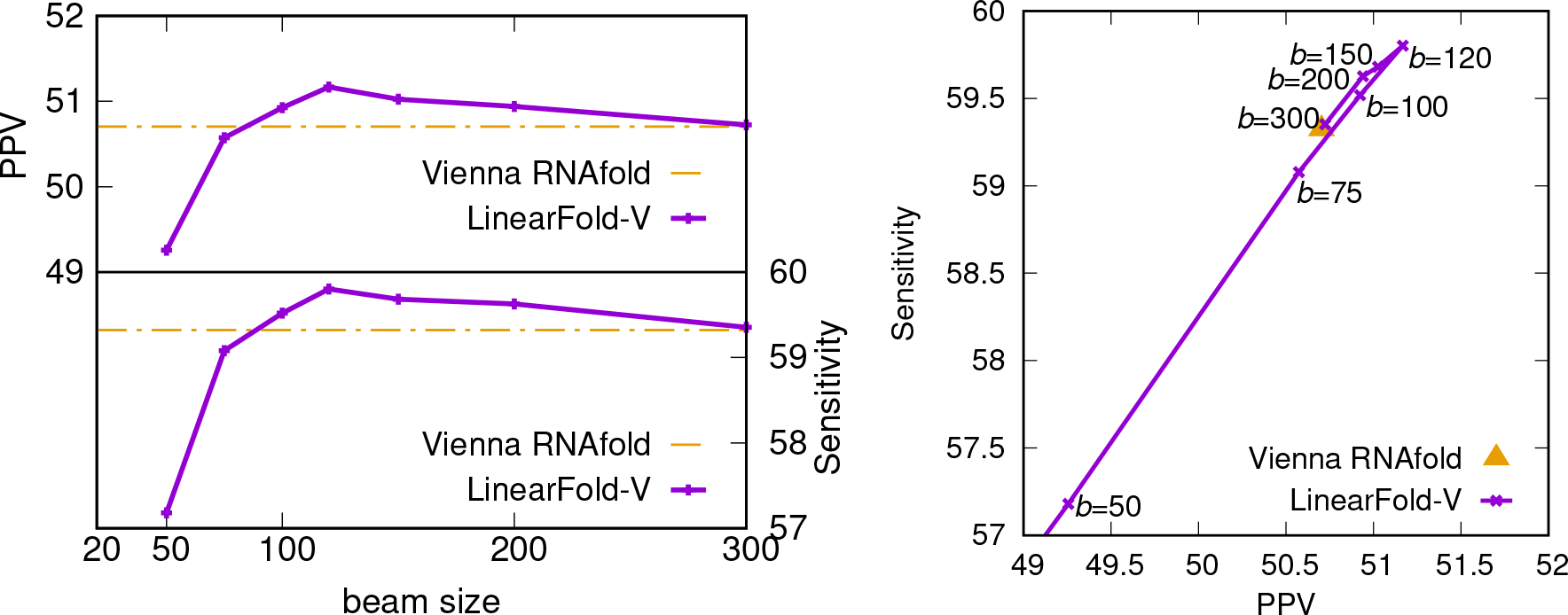
This figure corresponds to Figure 4(c)(d) but with the ViennaRNA version, running on the ArchiveII dataset. Left: trend of both PPV and Sensitivity with the increasing of beam size; right: PPV and Sensitivity of *LinearFold* by beam size.

**Fig. SI 2.**
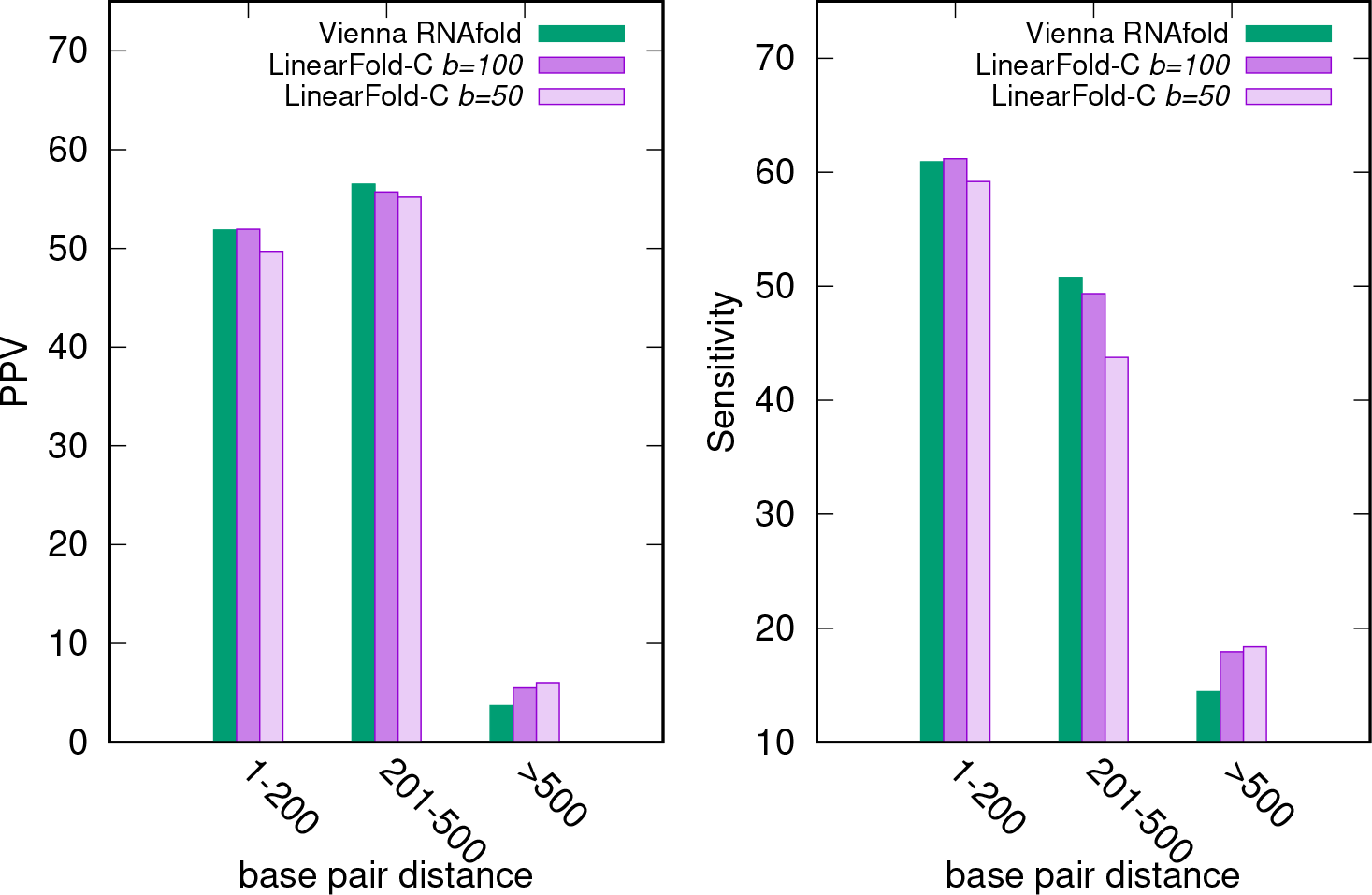
This figure corresponds to Figure4(c)(d), with the ViennaRNA version, running on the ArchiveII dataset. It shows PPV and Sensitivity of *LinearFold-V* by pair length in the Mathews dataset, comparing to Vienna RNAfold. Each bar represents the overall PPV/Sensitivity of all the base pairs in a length range.

**Fig. SI 3.**
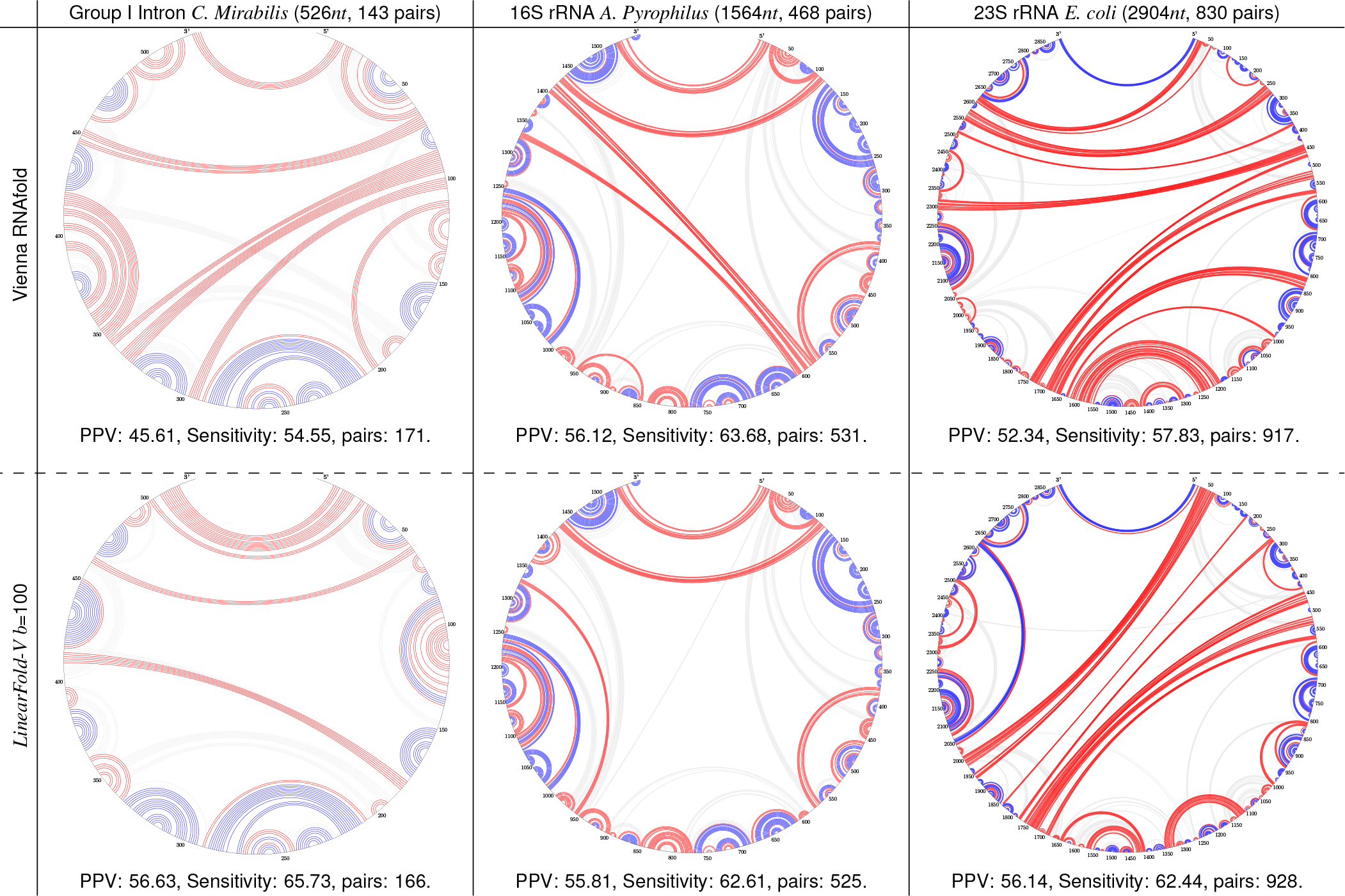
Circular plots of 3 RNA sequences (corresponding to Figure 5) comparing Vienna RNAfold with *LinearFold-V*.

**Fig. SI 4.**
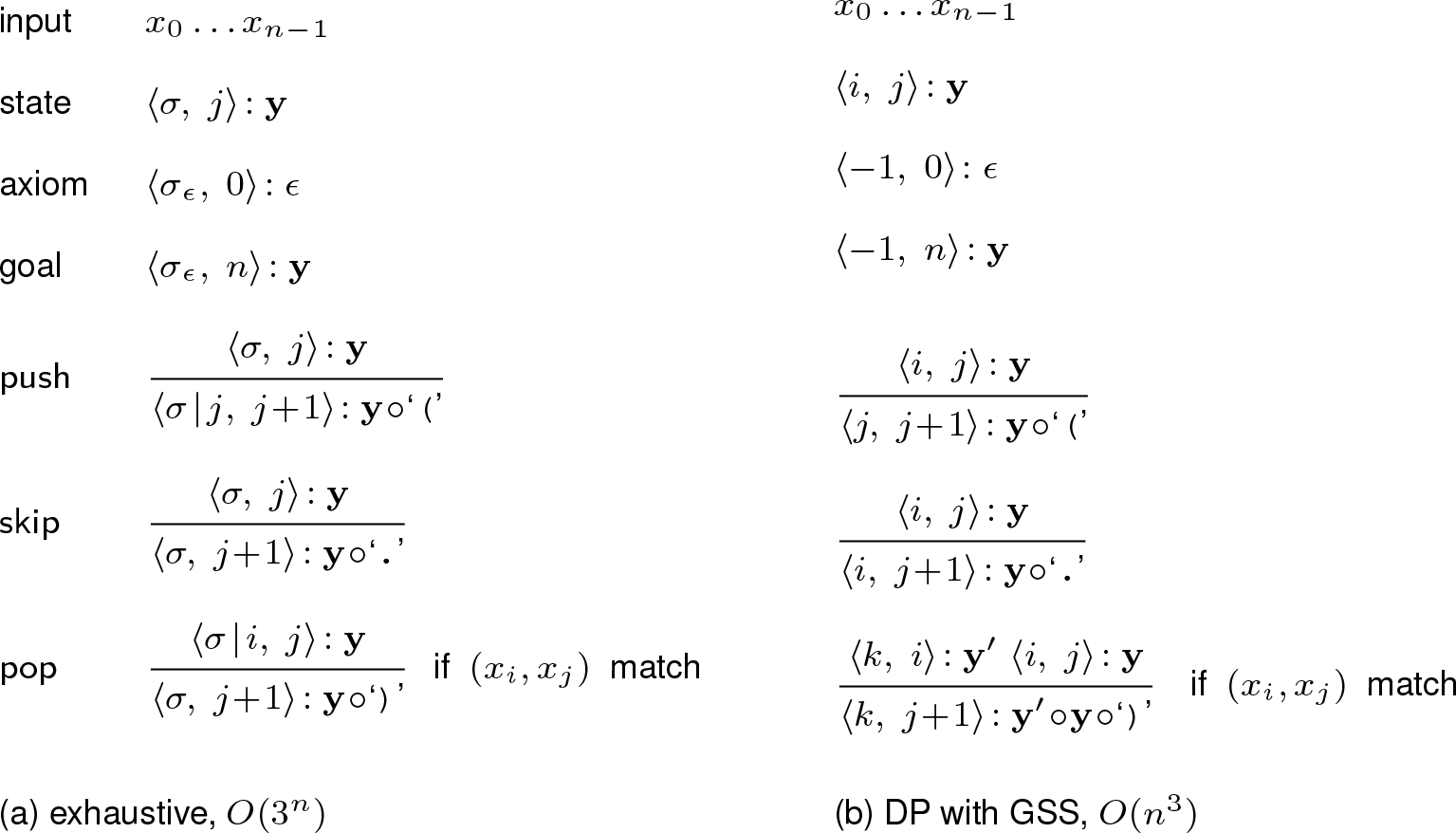
Deductive system comparing the exhaustive search algorithm and Dynamic Programming with Graph Structured Stack. Here o denotes string concatenation. We say (a, *b*) “matches” if (a, *b*) is one of the allowed pairs (C-G, A-U, G-U).

## Footnotes

* http://web.engr.oregonstate.eduHiukaib/demoJson+svg.html

Author contributions: L.H. conceived the idea based on D.H.’s suggestion. L.H., D.D., and K.Z. designed the algorithm. L.H. and D.D. implemented a prototype in Python. D.D. and K.Z. implemented the fast version In C++. D.H.M. and D.H. supervised testing of the algorithm. D.D. carried out testing and plotted figures. D.D., L.H., and D.H.M. wrote the manuscript; D.H. revised it.

